# Rho-mediated gene transcription promotes BRAF inhibitor resistance in de-differentiated melanoma cells

**DOI:** 10.1101/381806

**Authors:** SA Misek, KM Appleton, TS Dexheimer, EM Lisabeth, RS Lo, SD Larsen, KA Gallo, RR Neubig

**Affiliations:** Department of Physiology, Michigan State University, East Lansing, MI, 48824, USA; Department of Pharmacology, Michigan State University, East Lansing MI, 48824, USA; Division of Dermatology, Department of Medicine, University of California, Los Angeles, California, 90095, USA; Department of Molecular and Medical Pharmacology, University of California, Los Angeles, CA 90095, USA; Jonsson Comprehensive Cancer Center, University of California, Los Angeles, CA 90095, USA; David Geffen School of Medicine, University of California, Los Angeles, CA 90095, USA; Department of Medicinal Chemistry, University of Michigan, Ann Arbor, Michigan, 48109, USA; Vahlteich Medicinal Chemistry Core, University of Michigan, Ann Arbor, Michigan, 48109, USA

**Keywords:** BRAF inhibitor, resistance, RhoA, MRTF, YAP1, Melanoma

## Abstract

Over half of cutaneous melanoma tumors have BRAF^V600E/K^ mutations. Acquired resistance to BRAF inhibitors (BRAFi) remains a major hurdle in attaining durable therapeutic responses. In this study we demonstrate that approximately 50-60% of melanoma cell lines with vemurafenib resistance acquired *in vitro* show activation of RhoA family GTPases. In BRAFi-resistant melanoma cell lines and tumors, activation of RhoA is correlated with decreased expression of melanocyte lineage genes. Using a machine learning approach, we built gene expression-based models to predict drug sensitivity for 265 common anti-cancer compounds. We then projected these signatures onto the collection of TCGA cutaneous melanoma and found that poorly differentiated tumors were predicted to have increased sensitivity to multiple Rho kinase (ROCK) inhibitors. Two transcriptional effectors downstream of Rho, MRTF and YAP1, are activated in the Rho^High^ BRAFi-resistant cell lines, and resistant cells are more sensitive to inhibition of these transcriptional mechanisms. Taken together, these results support the concept of targeting Rho-regulated gene transcription pathways as a promising therapy approach to restore sensitivity to BRAFi-resistant tumors or as a combination therapy to prevent the onset of drug resistance.

## Introduction

Most cutaneous melanomas have BRAF^V600^ mutations ^12^. These mutations result in constitutive BRAF activity and downstream MAPK pathway activation, independent of upstream stimuli. The combination of BRAF inhibitors with MEK inhibitors was proposed as an approach to overcome BRAF inhibitor resistance ^29^ and it is clinically superior to BRAF inhibitor monotherapy against BRAF^V600^-mutant tumors ^2, 21, 35^. However, acquired resistance to the BRAF and MEK inhibitor combination is still common ^16^, consistent with non-MAPK pathway resistance mechanisms being important clinically ^14, 46^. Several resistance mechanisms have been identified which result in MAPK pathway reactivation, such as NRAS mutation or BRAF alternative splicing ^29, 34, 42, 43, 46, 51^. However, these MAPK-reactivating alterations only explain ∼50% of BRAFi-resistance mechanisms ^14, 16^. Some non-MAPK resistance mechanisms result from cancer cell intrinsic, epigenomically driven, adaptive responses to drug pressure early during therapy ^46^. These may result in wide-ranging phenotypic switches resulting in MAPK inhibitor resistance in patients and relapse ^14^. Melanoma cells grown without drug pressure stochastically switch between a rapid-cycling cell state and a rare slow-cycling cell state ^40^. Treatment with a BRAF inhibitor selects for cells in the slow cycling state ^43, 46^. These data are further supported by the observation that BRAFi/MEKi-resistant cells and tumors can be re-sensitized to treatment with BRAF or MEK inhibitors after a “drug holiday” ^4, 13, 25^.

The RhoA subfamily (RhoA, RhoB, and RhoC) of GTPases act as molecular switches which regulate actin dynamics. Herein, we will simply refer to the members of the RhoA subfamily generically as RhoA. The RhoA and RhoC isoforms are highly similar and often function redundantly in the cell, but in some contexts these two isoforms signal differently ^53^. In melanoma the RhoA subfamily, especially RhoC, promotes invasion and metastasis ^3, 18, 38^, and inhibition of the RhoA isoform suppresses tumor growth ^37^. Canonically, RhoA GTPases promote the formation of actin stress fibers by promoting G-actin polymerization and inhibiting F-actin depolymerization ^11, 45, 47^. Actin stress fibers have been shown to be increased in melanoma cells with acquired BRAFi resistance ^17^ and we confirm and extend that finding here.

In addition to regulating actin dynamics, RhoA GTPases also regulate gene transcription. This is, in part, through actin polymerization-dependent activation of MRTF and YAP1. MRTF and YAP1 are transcriptional co-activators which, upon activation, translocate into the nucleus and regulate gene transcription. Silencing of MRTF or SRF, a transcription factor by which MRTF modulates gene expression, prevents melanoma metastasis ^24^. Previously, we have developed a series of MRTF-pathway inhibitors including CCG-203971 and CCG-222740 ^1, 10, 15^ and demonstrated that CCG-203971 prevents melanoma metastasis, induces G1 cell cycle arrest, and reduces growth of melanoma cells ^10^. YAP1 has been shown to promote BRAFi/MEKi resistance in melanoma through suppression of apoptosis via BCL-xL and BIM dysgulation ^7, 14, 17, 22^; accumulation of YAP1 protein and enrichment of a YAP1 gene signature has been documented in about 40% of clinical melanoma samples from patients who relapsed on MAPK inhibitor therapies ^14^.

Previous studies have demonstrated that non-mutational, acquired resistance mechanisms represent a major hurdle in maintaining a durable response to MAPK-directed therapeutics ^14^. We hypothesize that activation of the RhoA pathway is one such acquired resistance mechanism. In this study we build upon existing literature to demonstrate that actin stress fiber accumulation and RhoA signaling are elevated in approximately half of vemurafenib-resistant melanoma cell lines tested and that this mechanism is also active in clinical tumors. RhoA^High^ but not RhoA^Low^-resistant lines are partially re-sensitized to vemurafenib by two structurally distinct ROCK inhibitors. We also demonstrate that RhoA activation is linked to loss of melanocyte lineage genes, a pattern also observed in human tumors. Finally, de-differentiated BRAFi-resistant cells have increased MRTF and YAP1 activation and these cells are more sensitive to pharmacological inhibition of these transcriptional mechanisms. De-differentiation of melanoma cells is a major mechanism of acquired BRAFi-resistance ^6, 32, 40, 48, 50^ and we have identified signaling alterations commonly associated with de-differentiation. This information is critical for developing therapeutic strategies to target this class of drug-resistant tumors.

## Materials and Methods

### Cell lines and culture

#### Selection of Vemurafenib-resistant cells

UACC62 and SK-Mel-19 cells were seeded into 10-cm tissue culture plates at ∼30% confluence and grown in DMEM as described below. After the cells had adhered to the plate (∼16 h), culture medium was supplemented with 2 µM vemurafenib. Medium was exchanged every 2-3 days for 10 mL of fresh media supplemented with 2 µM vemurafenib. Cells were split at a 1:3 ratio into a new 10-cm tissue culture plate when they reached ∼75% confluence (approximately 3-4 weeks) and approximately weekly for each subsequent passage. After two months of selection, cell populations were expanded in vemurafenib-containing media and frozen.

#### Sources of cells

Parental (P) and vemurafenib-resistant (R) M229P/R, M238P/R, and M249P/R cells were generously provided by Dr. Roger Lo at UCLA ^29^. SK-Mel-19 and UACC62 cells were obtained from Dr. Maria Soengas at The University of Michigan and were made resistant as described below.

Cells were cultured in DMEM (Gibco #11995-065) supplemented with 10% FBS (Gibco #10437-028) and 1% Antibiotic-Antimycotic (ThermoFisher, Waltham, MA, USA #15240062). Vemurafenib-resistant cells were continuously cultured in the presence of 2 µM vemurafenib. Cells were split at ∼75% confluence. Vemurafenib was removed from the culture medium when cells were seeded for experiments (e.g. immunofluorescence staining or qRT-PCR), except where otherwise indicated. Cells were routinely tested for mycoplasma contamination by DAPI staining. STR profiling on all cell lines was performed at the MSU genomics core. In all cases, isogenic pairs of cell lines had the same STR profile.

### Cloning

For CRISPR experiments the sgRNA sequences are listed in **(Table S1)**. These guide sequences were cloned into the pLentiCRISPRv2 vector (from Feng Zhang, Addgene plasmid #52961). All guide RNA sequences were confirmed by Sanger sequencing.

All cloning primers are listed in **(Table S1)**. Human RhoA^G12V^ was amplified and N-terminal HA-tagged. This PCR product was used as a template for a second stage of PCR amplification to add the Gateway adapter sequences. Human MRTFA was amplified out of the p3xFLAG-MRTFA vector (Addgene plasmid#11978) and tagged with gateway adapters which preserve the N-terminal 3x FLAG tag from the vector. The RhoA and MRTFA PCR products were first cloned into pDONR221 using the Gateway BP Clonase II Enzyme Mix from ThermoFisher (#11789020) using the manufacturer’s protocol. RhoA, MRTFA, and Gus (which is included in the BP reaction kit) were subcloned into the pLX301 vector (from David Root, Addgene plasmid #25895) using the Gateway LR Clonase II Enzyme mix from ThermoFisher (#11791020). The presence of the correct insert in the final plasmid was confirmed by Sanger sequencing.

### Virus Preparation and Infection

HEK-293T cells were seeded into 10-cm plates and were allowed to attach overnight. The next day, the cells were transfected with a plasmid cocktail containing 5000 ng of the pLentiCRISPRv2 or pLX301 plasmid, 3750 ng of psPAX2, 1250 ng of pMD2.G, and 20 uL of Lipofectamine 2000 in 400 uL of OptiMEM. The next morning the media was changed to 10 mL of fresh culture medium, and the next day each plate was supplemented with an additional 5 mL of culture medium. After 24 h, the culture medium was harvested and filtered through a 0.45-micron syringe filter. Virus was stored at 4 C and was used within 2 weeks.

UACC62P and UACC62R cells were seeded into 10-cm plates and were allowed to attach overnight. The next day the medium was changed to 10-mL of complete medium and 1 mL of virus supernatant was added in the afternoon. The next morning the medium was changed and the cells were incubated an additional 24 h then the cells were treated with 10 µg/mL puromycin until all of the control cells died (approximately 72 h). For all virus experiments, the cells were used within 1-2 passages and each biological replicate for each experiment used a different batch of cells. We did not pick individual clones for the CRISPR cell lines, but instead used a pooled infection approach. Validation of CRISPR knockout efficiency was done by immunoblotting for the target protein.

### Compounds and Antibodies

Vemurafenib (#S1267), Y-27632 (#S1049), fasudil (#S1573), and dasatinib (#S1021) were purchased from Sellekchem, Houston, TX, USA. CCG-222740 ^15^ was synthesized in the lab of Dr. Scott Larsen at the University of Michigan. All compounds were diluted in DMSO to a stock concentration of 10 mM. Compounds were frozen at −20 °C. Antibodies against YAP1 (#14074), MLC2 (#3672), pMLC2 (#3674), and Sox10 (#89356) were purchased from Cell Signaling, Danvers, MA, USA. Antibodies against MRTF-A (#sc21558), MRTF-B (#sc98989), and Actin (#sc1616) were purchased from Santa Cruz, Dallas, TX, USA. Donkey anti-Mouse800 (#926-32212), Donkey anti-Goat680 (#926-68074), and Donkey anti-Rabbit680 (#926-68073) immunoblotting secondary antibodies were purchased from LI-COR, Lincoln, NE, USA. Alexa Fluor goat anti-rabbit488 (#A11034) and donkey anti-goat488 (#A11055) were purchased from Invitrogen. Alexa Fluor546 Phalloidin (#A22263) was purchased from ThermoFisher.

### qRT-PCR

Cells were cultured and treated as indicated, rinsed once in PBS, and total cellular RNA was harvested with the Qiagen, Hilden, Germany, RNeasy kit (#74104) according to the manufacturer’s protocol. RNA was eluted in nuclease-free H_2_O. cDNA was synthesized using the High-Capacity cDNA RT kit from ThermoFisher (#4368814) from 1000 ng of total RNA, according to the manufacturer’s protocol. qPCR was performed using the SYBR Green PCR Master Mix (#4309155) from ThermoFisher according to the manufacturer’s protocol using an Agilent Mx3000P qPCR instrument. Primers were purchased from Integrated DNA Technologies, San Jose, CA, USA and the primer sequences are included in **(Table S1)**. Primers were designed using the Harvard Primer Bank tool (https://pga.mgh.harvard.edu/primerbank/). Fold-change analysis was performed using the ΔΔCT method.

### RNA-Seq sample preparation and data processing

Total cellular RNA was extracted from UACC62P and UACC62R cells (two biological replicates per cell line) using the same method which was used for qPCR experiments. RNA concentration was measured by Qubit and quality control was performed on an Agilent 2100 Bioanalyzer in the MSU Genomics Core. All RNA samples had a RIN score > 8. Barcoded libraries were prepared using the Illumina TruSeq Stranded mRNA Library Preparation Kit on a Perkin Elmer Sciclone G3 robot following manufacturer’s recommendations. Completed libraries were QC’d and quantified using a combination of Qubit dsDNA HS and Caliper LabChipGX HS DNA assays. Libraries were pooled and run on two lanes, and sequencing was performed in a 1×50 bp single-end read format using HiSeq 4000 SBS reagents. Base calling was done by Illumina Real Time Analysis, RTA_ v2.7.7 and output of RTA was demultiplexed and converted to FastQ format with Illumina Bcl2fastq v2.19.0. Sequencing was performed at a depth of >30M reads/sample. Quality control was performed on the FastQ files using FastQC v0.11.5, and reads were trimmed using Trimmomatic v0.33. Reads were mapped using HISAT2 v2.1.0 and analyzed using HTSeq v0.6.1. Differential gene expression was calculated using edgeR. Raw RNA-Seq reads and processed HTSeq read counts are available on GEO under GSE115938.

### Immunoblotting

Cells were cultured and treated as indicated, placed on ice, and rinsed once in cold PBS. Cells were lysed in 2x Laemmli Sample Buffer (Biorad, #1610737). Samples were sonicated with a probe sonicator for approximately 5 sec, then boiled at 100 °C for 10 min. Samples were loaded onto a 12% polyacrylamide gel and transferred to Immobilon-FL PVDF Membrane (Millipore Sigma, Burlington, MA, USA, #IPFL00010). Membranes were blocked in 5% BSA + TBS-Tween (1:1000) for 1 h, then incubated in primary antibody overnight at 4 °C. Membranes were washed 3x in TBS-Tween and were then incubated in the appropriate secondary antibody at a 1:20,000 dilution for 1 h at room temperature. All antibodies were diluted in blocking buffer. Membranes were washed 3x in TBS-Tween then dried and imaged on a LI-COR Odyssey FC imaging system.

### Immunofluorescence staining

Cells were seeded into 8-well chamber slides and were treated as indicated in the figure legends. Cells were fixed with 3.7% formaldehyde for 15 min then blocked in 2% BSA PBS-Triton (0.1%) for 1 h at room temperature. Cells were incubated overnight at 4 °C in primary antibody at a 1:100 (MRTF-A or MRTF-B) or 1:500 (YAP1) dilution in blocking buffer. Cells were washed 3x in PBS then were incubated in the appropriate secondary antibody at a 1:1000 dilution for 1 h at room temperature. Cells were washed 3x in PBS then were mounted in ProLong Gold Antifade + DAPI (ThermoFisher, #P36935). Slides were cured overnight at room temperature and were then imaged on a Nikon TE2000-U Fluorescence Microscope at 20x magnification.

Cells were stained with Alexa Fluor546 Phalloidin (#A22263) to visualize F-Actin. For these experiments, cells were fixed and blocked as described above. Cells were then incubated in Phalloidin diluted 1:100 in blocking buffer for 1 h at room temperature before being washed and mounted. For all immunofluorescence experiments, images were blinded by an independent party or an automated R script before quantification. For a cell to be considered as stress fiber-positive, the cell was required to contain at least one stress fiber which spanned >90% the length of the cell. We repeated all staining experiments at least 3 times and typically analyzed at least 10 fields per biological replicate. In total we analyzed at least 400 cells per experimental group, but in most cases over 1000 cells per experimental group. For subcellular localization experiments, data are represented as a stacked bar graph wherein the fraction of cells that have predominantly nuclear, pan-cellular, or cytosolic localization is plotted as a fraction of the total cells. A cell was considered to have “cytosolic” localization if there was clear nuclear exclusion. Inversely a cell was described as having “nuclear” localization if the staining intensity was appreciably higher than in the cytosol. If there was no apparently difference between the nuclear and cytosolic staining then the cell was described as having “pan-cellular” distribution of the protein being assessed.

### Cell viability experiments

Cells were seeded into 384-well tissue culture plates (PerkinElmer, Waltham, MA, USA, #6007689) at a density of 1000 cells/well in 20 uL of media and were allowed to attach overnight. The next day, drugs were pre-diluted at 4x final concentration in culture medium then added to the 384-well plates so that the final volume was 40 µL/well. For the single compound dose response experiments, the compound was pre-diluted at 2x the final concentration and 20 uL was added to each well. A PBS barrier was added to the outer wells of the plate to limit evaporation. Cells were cultured under these conditions for 72 h. To assess viability, 10 µL of CellTiter-Glo (Promega, Madison, WI, USA, #G7573) was added to each well. Plates were incubated for 5 min at room temperature then briefly centrifuged (4000 rpm, 60 seconds) before being read on a Bio-Tek Synergy Neo plate reader. Data are plotted versus Vemurafenib concentration for each treatment condition. The Area Under the Curve (AUC) was calculated for each curve using GraphPad Prism.

### Bioinformatics

#### Dataset Processing

Cancer Cell Line Encyclopedia (CCLE) gene expression Affymetrix CEL files (Version 19-Mar-2013) were downloaded from the Broad Institute CCLE data portal. CEL files were processed using Affymetrix Expression Console (Build 1.4.0.38). Probe IDs were collapsed to gene names using the CollapseDataset function on GenePattern. The TCGA RNA-Seq dataset for cutaneous melanoma (SKCM) was downloaded from the UCSC Cancer Genome Browser portal. No further data processing was performed prior to analysis.

RNA-Seq data for 62 human tumors paired for pre- and post-MAPK inhibitor resistance was downloaded from GSE65185 ^14^. Analysis of these data was performed on the pre-processed CuffnormFPKM dataset included in this series. RNA-Seq data for *in vitro* generated vemurafenib-resistant M229P/R and M238P/R cells was downloaded from GSE75313 ^46^. These data were processed using the above described RNA-Seq data processing pipeline. Melanoma ssRNA-Seq data was downloaded from GSE72056 and filtered to include only melanoma cells. Missing values were imputed with the MAGIC algorithm ^52^.

Data for the M229 cells treated with vemurafenib for different times was downloaded from GSE110054. No further processing was performed on this dataset prior to ssGSEA analysis.

#### Gene Ontology/KEGG pathway analysis

Using the CCLE dataset, 38 adherent cell lines with BRAF^V600^-mutations were identified. For all cell lines, PLX4720 (activity area) was correlated with gene expression. Genes highly expressed in resistant cells (genes with a Pearson correlation coefficient < −0.5 when correlated with PLX4720 sensitivity) and genes weakly expressed in resistant cells (Pearson correlation coefficient > 0.5) were identified. Gene ontology and KEGG pathway analysis was performed on the gene sets using GATHER (http://changlab.uth.tmc.edu/gather/gather.py) with network inference.

#### GSEA/ssGSEA

GSEA (v19.0.24) was performed using GenePattern (http://software.broadinstitute.org/cancer/software/genepattern/) with ‘number of permutations’ = 1000, and ‘permutation type’ = phenotype. All other parameters were left as default. ssGSEA (9.0.9) was performed on GenePattern with all parameters left as default. The ssGSEA output values were z-score normalized.

A RhoA/C gene signature was generated by using all genes which are upregulated > 2-fold by overexpression of either RhoA or RhoC from the GSE5913 dataset in NIH-3T3 cells. These two lists were merged, and duplicates were removed. This resulted in a list of 79 genes (Supplemental Table 1).

The melanocyte lineage signature included all genes in the GO_MELANIN_METABOLIC_PROCESS (GO: 0006582) and GO_MELANOCYTE_DIFFERENTIATION (GO: 0030318) MSigDB signatures. The combined list was filtered to remove duplicate genes.

The YAP1 signature used was the CORDENONSI_YAP_CONSERVED_SIGNATURE in the C6 collection on MSigDB. The MRTF signature is comprised of all genes downregulated > 2-fold upon MRTF knockdown in B16F2 melanoma cells ^24^ (Supplemental Table 1).

#### Drug Response Signatures

The correlated gene expression profiling and drug IC50 values were downloaded from the GDSC data portal (https://www.cancerrxgene.org/downloads). Gene expression data was median centered so that the median expression of each gene across the cell lines was equal to 0. Data was randomly divided into a training (80%) and test (20%) set. A predictive model was built on the training set for each compound (n = 265 compounds) using a random forest algorithm (randomForest package in R) with ntrees = 500 and mtry = sqrt(#genes). Each model was validated on the test dataset by calculating the Pearson correlation coefficient between the predicted and actual IC50s. Models with a Pearson correlation coefficient > 0.3 were considered predictive. A full table of these results is included as **(Table S2)**. To use gene expression data to predict drug response on clinical tumors, the TCGA SKCM data were median-centered using the same method used on the GDSC training data. Since the TCGA and GDSC datasets were collected on different gene expression analysis platforms, the two datasets were filtered to include only overlapping genes. Models from GDSC which were deemed predictive for a drug response were then projected onto the TCGA dataset. Melanocyte Lineage signature scores of TCGA samples **(Fig. 2D)** were negatively skewed from a normal distribution (corrected z^3^ = − 1.94). Of the 473 tumors, 70 were > 2 SD below the mean and none > 2SD above the meal. Consequently, samples at least 2 SD below the mean are considered “lineage low” and samples greater than 2 SD below the mean are considered “lineage high”. The average predicted IC_50_ for the Lineage low and Lineage high tumors was calculated by averaging the predicted log(IC_50_) for each sample class.

**Figure 1.**
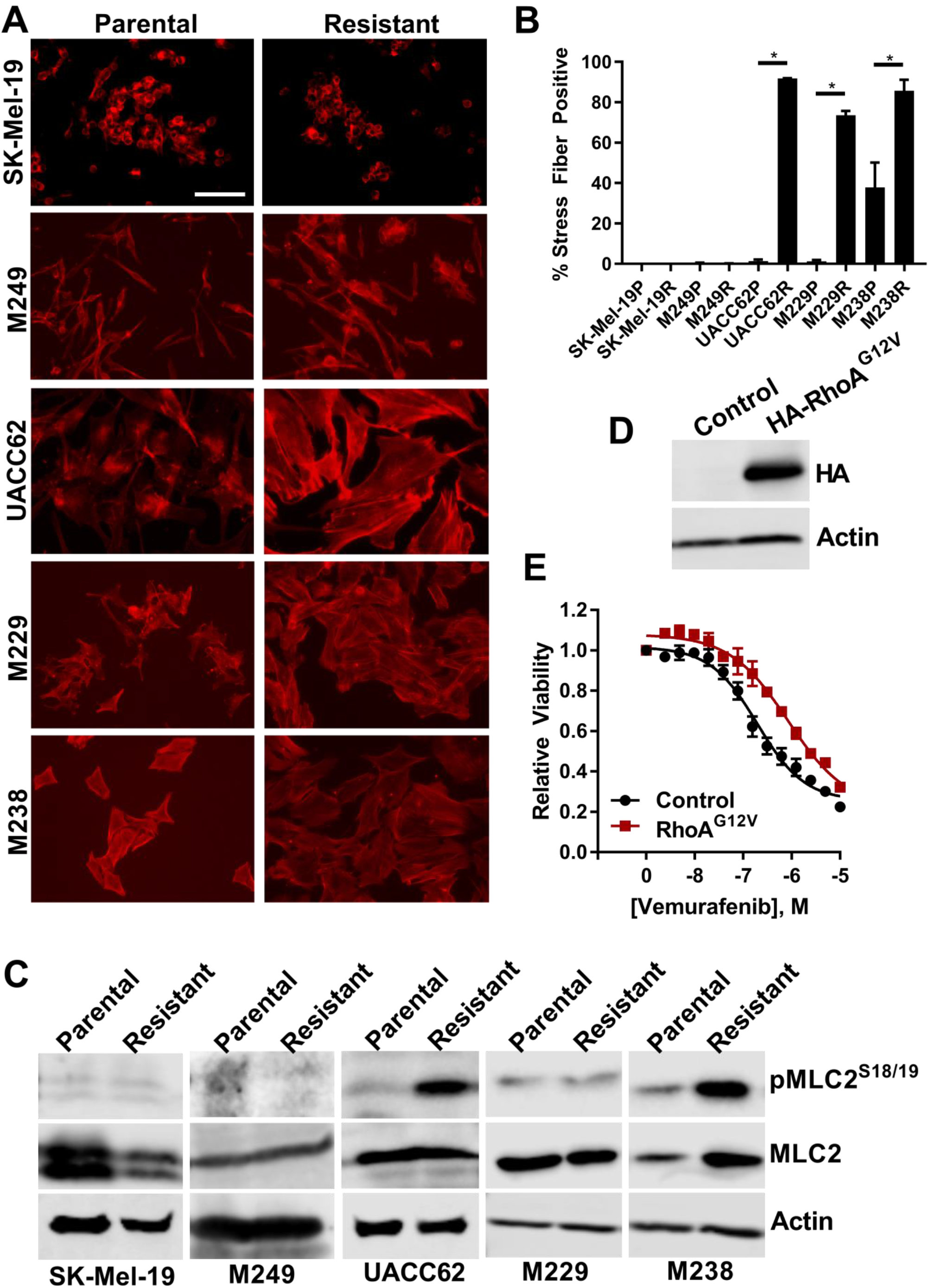
RhoA is activated in BRAFi-resistant melanoma cells and tumors. **A)** Cells were seeded into 8-well chamber slides and were allowed to attach overnight. The next day cells were fixed and stained with fluorescently labeled phalloidin. Representative images from n = 3 biological replicates and n = 1 technical replicate. Scale bar is 10 µm. **B)** Actin stress fiber positive cells were quantified using ImageJ. Statistical analysis was performed using unpaired t-tests to compare matched parental and resistant lines. * indicates that p < 0.05. **C)** MLC2^S18/19^ phosphorylation in UACC62 (left) and SK-Mel-19 (right) parental and resistant cells was assessed by immunoblotting. Total MLC2 and Actin were used as loading controls. Representative blots from n = 3 biological replicates and n = 1 technical replicate. **D)** UACC62P cells stably expressing HA-RhoA^G12V^ were lysed and immunoblotted with anti-HA and anti-Actin antibodies. Representative images from n = 3 biological replicates and n = 1 technical replicate. **E)** UACC62P cells stably expressing Gus (control) or HA-RhoA^G12V^ were seeded into 384-well plates and treated with a 14-point vemurafenib concentration gradient with a top dose of 10 µM as described in the materials and methods. Data is average from n = 3 biological replicates with n = 3 technical replicates.

**Figure 2.**
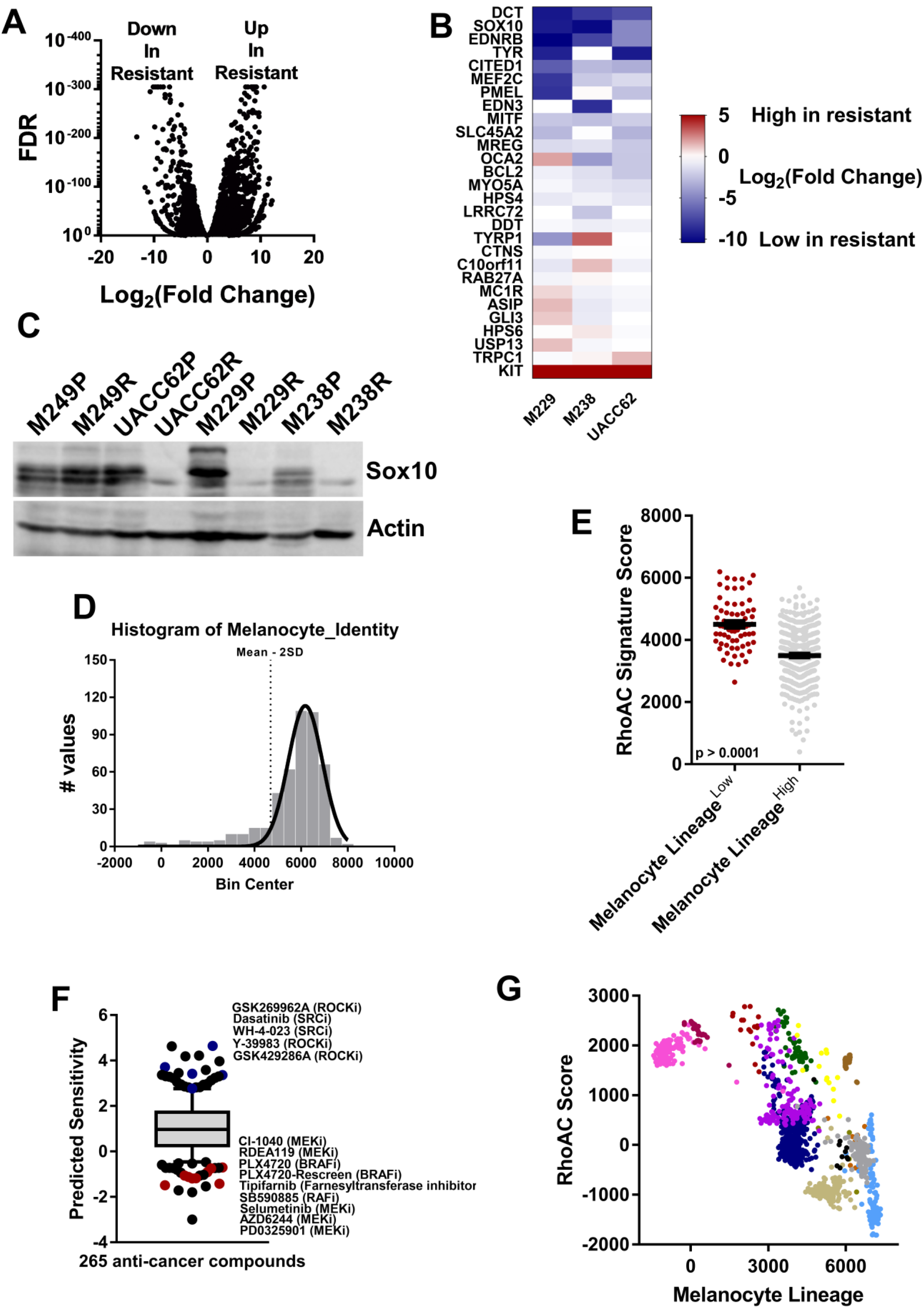
Melanoma differentiation status is inversely correlated with Rho activation. **A)** RNA-Seq was performed on parental (UACC62P) and Vemurafenib-resistant (UACC62R) cells. Differential gene expression was visualized on a volcano plot. n = 2 biological replicates and n = 1 technical replicate per treatment condition. **B)** Heatmap of differential expression of Melanocyte Lineage signature genes in M229P/R, M238P/R, and UACC62P/R cells. Blue indicates that the gene is downregulated in the resistant cell line, and red indicates that the gene is upregulated in the resistant cell line. **C)** Sox10 protein expression was assessed across a panel of 4 parental and resistant melanoma cell lines. Actin was used as a loading control. Representative image from n = 3 biological replicates and n = 1 technical replicats. **D)** Histogram of Melanocyte Lineage signature scores for samples in the SKCM TCGA dataset (n = 473). Dotted line represents 2 SD below the mean of the Gaussian fit. Samples were stratified into Melanocyte Lineage high and Melanocyte lineage low samples as described in the Materials and Methods section. **E)** RhoA/C activation was predicted on TCGA SKCM tumors using ssGSEA with a RhoA/C gene signature. Tumors were stratified based on expression of the Melanocyte Lineage signature genes as described in Materials and Methods. **F)** Drug signatures for 265 common anti-cancer drugs in the GDSC dataset were generated using a random forest algorithm. These signatures were projected onto the TCGA SKCM RNA-Seq dataset and predicted drug sensitivity for each tumor was calculated. Tumors were stratified by expression of Melanocyte Lineage signature genes as described in Materials and Methods. **G)** Correlation of the RhoA/C signature and the Melanocyte Lineage signature in a single cell RNA-Seq dataset ^49^ from melanoma cells isolated from human tumors. Different colors represent different tumors of origin for the cell line populations.

### Statistical Analysis

Most bioinformatics analysis was performed using R v3.3.0. Data analysis and statistics were performed using GraphPad Prism v6 or v7. Dose response curves were fit using nonlinear least square regression [log(agonist) vs. response – Variable slope (four parameters)]. The AUC was calculated for each dose response curve in GraphPad Prism over a vemurafenib concentration range of 10^−9^ to 10^−5^. Datasets with two samples per group were analyzed by unpaired two-tailed t-tests. Pearson correlation coefficients were calculated in R (for drug response signatures) or GraphPad Prism (for all other analysis). Data are presented as mean ± S.E.M, and a p-value < 0.05 was considered statistically significant.

## Results

### RhoA activation in BRAFi-resistant melanoma cells and tumors

We analyzed a panel of matched parental and BRAFi-resistant melanoma cell lines and found that three of the resistant cell lines (UACC62R, M229R, and M238R) assumed a fibroblast-like morphology, while there was no overt change in the other two resistant cell lines (SK-Mel-19R and M249R). Since cell shape is largely controlled through modulation of the actin cytoskeleton, we examined F-actin structure by staining the cells with fluorescently labeled phalloidin. There was an increase in the number of actin stress fiber-positive UACC62R, M229R, and M238R cells compared to matched parental control cell lines, but there was no overt change in stress fiber levels in the SK-Mel-19R and M249R cells **(Fig. 1A/B)**. Since an increase in stress fibers would suggest that Rho activation is altered, we also analyzed MLC2 phosphorylation in the matched parental and resistant cell lines. MLC2 phosphorylation is increased in the UACC62R and M238R cell lines, but not in SK-Mel-19R or M249R cells **(Fig. 1C)**. Interestingly, there was no change in MLC2 phosphorylation in the M229R cells despite the fact that they are stress fiber positive, which may suggest that these cells utilize an alternative signaling mechanism to activate RhoA and increase stress fibers.

These data suggested that RhoA was activated in the resistant cell lines, but it was not clear whether RhoA itself was functionally important in BRAFi resistance. To address this question we generated UACC62P cells which stably express RhoA^G12V^ **(Fig. 1D)**. This specific mutation is not found in any SKCM tumors in the TCGA dataset, however, the constitutively active RhoA^G12V^ model is a useful tool for studying mechanisms of Rho signaling since it is independent of upstream stimuli. Consistent with our observations suggesting that RhoA is activated in a subset of the resistant cell lines, overexpression of RhoA^G12V^ reduced vemurafenib sensitivity by approximately 8-fold **(Fig. 1E)**.

To more broadly confirm this finding, we correlated cell sensitivity to PLX4720 (a BRAF inhibitor which is structurally similar to vemurafenib) with the gene expression results for 38 BRAF^V600^-mutant cell lines from the Cancer Cell Line Encyclopedia (CCLE). Genes which are highly expressed in PLX4720-resistant cells (genes with a Pearson correlation of gene expression values vs drug activity area < −0.5) were analyzed by Gene Ontology and KEGG pathway analysis. One of the most statistically significant GO terms was “small GTPase mediated signal transduction” **(Fig. S1)** and the most statistically significant KEGG pathway was “Regulation of actin cytoskeleton” **(Fig. S1)**. A RhoA/C gene signature was also inversely correlated (R = −0.42) with PLX4720 sensitivity **(Fig. S2)**. Collectively, these data support the idea that RhoA activation is positively correlated with BRAFi resistance across a wide array of melanoma cell lines. To determine whether these cell line observations are applicable in the clinical context, we analyzed RNA-seq data from 41 tumors before and after development of resistance to BRAFi/MEKi ^14^. More than half of the resistant tumors (n = 24) had an increased RhoA/C signature score over the baseline tumor **(Fig. S3)**. Taken together, these data suggest that RhoA is activated in approximately half of BRAFi-resistant cells and tumors, and that RhoA activation is inversely correlated with BRAFi sensitivity.

Since the most common class of BRAFi resistance mechanisms is through MAPK re-activation we then wondered whether RhoA activation was mutually exclusive with MAPK reactivation-mediated resistance. If resistance is developed through MAPK re-activation then the resistant cells should retain ERK phosphorylation when treated with vemurafenib. As expected, vemurafenib inhibits ERK phosphorylation in all 5 parental cell lines. Vemurafenib fails to inhibit ERK phosphorylation in the two RhoA^Low^ resistant lines (SK-Mel-19R and M249R), which in the case of M249R is expected since these cells develop resistance by developing an NRAS^Q61K^ mutation. In the three RhoA^High^ resistant cell lines, vemurafenib partially inhibited ERK phosphorylation (M229R and UACC62R) but failed to suppress ERK phosphorylation in M238R cells **(Fig. S4)**. This finding is important since it suggests that Rho may be important even in cells which harbor MAPK-reactivating resistance mechanisms.

### Resistant cell lines with a low level of melanocyte differentiation show high RhoA activity

We next wanted to understand mechanistically why the RhoA pathway is only activated in a subset of vemurafenib-resistant cells. We performed RNA-Seq on the UACC62P/R cells **(Fig. 2A)**, and also analyzed published RNA-Seq data for the M229P/R and M238P/R cells. The most striking finding was that a number of genes linked to the melanocyte lineage and pigment production were downregulated in all three of the RhoA^High^ resistant cell lines. To more quantitively analyze this phenotype we generated a “Melanocyte Lineage” gene signature **(Table S3)** which is comprised of genes involved in pigment production and the melanocyte lineage. A majority of the signature genes are downregulated in all three of the Rho^High^ resistant cell lines **(Fig. 2B)** which suggests that loss of melanocyte identity is associated with Rho activation in BRAFi-resistant cells. There is also a temporal association between expression of the melanocyte lineage genes and RhoA/C signature genes **(Fig. S5)**. One of the most strongly downregulated genes, at the mRNA level, is the transcription factor Sox10 which is one of the “master regulators” of the melanocyte lineage; we confirmed that Sox10 is also downregulated at the protein level **(Fig. 2C)**. Interestingly, there was no change in Sox10 expression in the M249P/R cells which did not have increased stress fibers **(Fig. 2C)**. We also found that Sox9 is upregulated at the mRNA level in all three of the RhoA^High^ resistant cell lines but not in the RhoA^Low^ resistant lines **(Fig. S6)**. These results are consistent with previous findings which suggest that Sox10 suppresses Sox9 expression ^41^, and suggest that this switch in transcription factor expression may be reflective of the differentiation status of the resistant cells.

We next wanted to determine whether this de-differentiation phenotype was also important in human SKCM tumors. We projected the “Melanocyte Lineage” signature onto the SKCM TCGA dataset and then fit a Gaussian distribution to the signature scores but the distribution was skewed towards lower signature scores (corrected z^3^ = −1.94). Most of the tumors fell within 2 standard deviations of the mean, but there was a subset of tumors (n = 70) which had low expression of melanocyte lineage genes (low was defined at being > 2SD below the mean) **(Fig. 2D)**. There were no tumors which had a signature score > 2 SD above the mean. As expected, tumor purity was correlated with the expression of melanocyte lineage genes **(Fig. S7)**, but this does not fully explain why these tumors have lower expression of these genes given the magnitude of the downregulation of the melanocyte lineage signature. Consistent with the finding that RhoA is activated in de-differentiated BRAFi-resistant cell lines, we also found that tumors with decreased expression of melanocyte lineage genes have increased expression of RhoA/C target genes **(Fig 2E)**.

The small number of tumors (n = 70 out of 473 total tumors) which have decreased expression of melanocyte lineage genes may be due to the fact that all of the tumors in this dataset were treatment-naïve with respect to BRAF inhibitors. But since the transcriptional profile of these lineage-low tumors is similar to that of the BRAFi-resistant cell lines, it is possible that these tumors may have intrinsic resistance to BRAF inhibitors. To test this hypothesis we generated gene expression signatures using a random forest machine learning algorithm to predict drug response for 265 common anti-cancer drugs. These signatures were then projected onto the TCGA dataset to predict drugs to which the de-differentiated tumors are differentially sensitive **(Fig. 2F, Table S4)**. As expected, the de-differentiated tumors are predicted to be less sensitive to multiple BRAF and MEK inhibitors, including PLX4720 (a structurally similar vemurafenib analog). These predictions support the idea that melanoma tumors with a de-differentiated transcriptional signature are less sensitive to BRAF inhibition even before selection by BRAFi treatment. This supports what we observed in experimentally derived resistant cell line models. Also, de-differentiated tumors are predicted to have increased sensitivity to multiple ROCK inhibitors which is interesting since ROCK is one of the canonical RhoA effector proteins ^9, 28^. This again correlates with the sensitivity of our three resistant lines to ROCK inhibitors.

The observation that RhoA activation is inversely correlated with differentiation status in human tumors could be marred by the contribution of non-malignant cells to the overall bulk gene expression profile of the tumor. For example, it is expected that in some cases cancer-associated fibroblasts or endothelial cells might have high RhoA activity ^31, 55^. To more directly address the hypothesis that differentiation status is inversely correlated with Rho activation in *melanoma cells* we used publicly available single cell RNA-Seq data ^49^ to correlate a RhoA/C signature and the Melanocyte Lineage signature. As expected, cells clustered together based on their tumor of origin which is due to the strong inter-tumor transcriptomic heterogeneity ^49^. Even within a single tumor, poorly differentiated cells have elevated RhoA activation **(Fig. 2G)**. In total, these data suggest that tumors which acquire a de-differentiated phenotype have elevated RhoA activation and are predicted to be more sensitive to inhibition of RhoA signaling.

### ROCK inhibition sensitizes RhoA^High^ BRAFi-resistant melanoma cells

It is difficult to pharmacologically target RhoA directly, so an alternative approach is to target downstream effector pathways. Since we predicted that poorly differentiated human melanoma tumors are more sensitive to ROCK inhibitors it is possible that de-differentiated BRAFi-resistant cells are more sensitive to ROCK inhibitors. It is also possible that ROCK inhibition might re-sensitize the resistant cells to Vemurafenib. To test this hypothesis we used two ROCK inhibitors, Y-27623 and Fasudil, which have structurally distinct chemical scaffolds. RhoA^High^ BRAFi-resistant cells (but not RhoA^Low^ resistant cells) are more sensitive to either of the ROCK inhibitors as a single agent **(Fig. S8)**. ROCK inhibition also re-sensitizes RhoA^High^ (but not RhoA^Low^) BRAFi-resistant cells to vemurafenib **(Fig. 3A-H)**. Re-sensitization to vemurafenib was most pronounced in M229R cells **(Fig. 3I-K)** which is interesting since these cells do not have increased MLC2 phosphorylation. Since increased sensitivity to ROCK inhibitors alone, or the effect of ROCK inhibitors on re-sensitizing cells to vemurafenib is only observed in cells which have increased stress fibers it suggests that this combination treatment may be specific for cells/tumors which activate this signaling mechanism.

**Figure 3.**
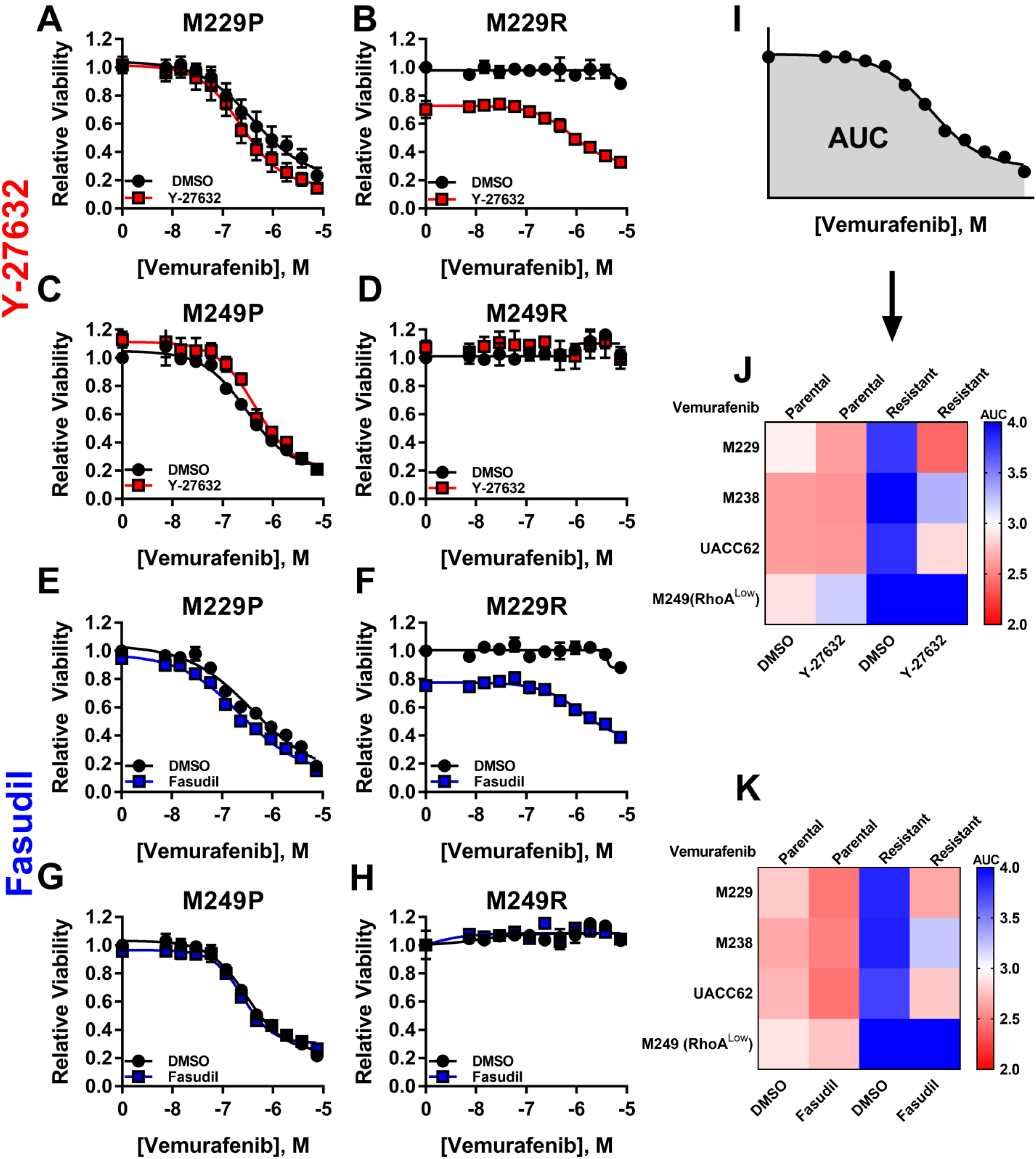
ROCK inhibition reverses BRAFi resistance in RhoA^High^ BRAFi-resistant melanoma cells. Parental and Vemurafenib-resistant cell lines were seeded into 384-well plates at a density of 1,000 cells/well and cells were allowed to attach overnight. The next day, cells were treated with Vemurafenib at the indicated concentrations with or without the ROCK inhibitors Y-27632 (red) or Fasudil (blue) at 10 µM. Cells were grown for 72 h then viability was measured with CellTiter-Glo. Pooled viability data from n = 3 biological replicates and n = 1 technical replicates. **A-H)** Cell lines were treated as labeled with ROCK inhibitors (Y-27632 or Fasudil) along with Vemurafenib. **I)** Schematic of Area Under Curve (AUC) calculation. Larger AUC indicates lower sensitivity to the drug combination and smaller AUC indicates greater sensitivity to the drug combination. **J)** Heatmap of AUC values for the Vemurafenib/Y-27632 drug combination for four parental and resistant cell line pairs. **K)** Heatmap of AUC values for the Vemurafenib/Fasudil drug combination for four parental and resistant cell line pairs.

### MRTF and YAP activation in RhoA^High^ BRAFi-resistant cells

In addition to modulating cytoskeletal re-arrangement, RhoA also regulates gene expression. Two transcriptional co-activators downstream of RhoA are YAP1 and MRTF. MRTF and YAP1 have similar transcriptional outputs and can perform redundant functions in several contexts ^8, 56^. To determine whether YAP1 and MRTF are activated in RhoA^High^ BRAFi-resistant cells, we measured the subcellular localization of YAP1 and MRTF-A **(Fig. 4A/B)**. YAP1 nuclear localization is elevated in M229R and M238R cells compared to matched parental cell lines and is elevated to a lesser extent in UACC62R cells. The converse is true with respect to MRTF-A localization since nuclear MRTF-A is increased in UACC62R cells but not M229R or M238R cells **(Fig. 4A/B)**. Several YAP1- and MRTF-related genes are highly expressed BRAF-mutant cell lines with intrinsic BRAFi resistance **(Fig. S9)**. These include the YAP1/MRTF target gene CYR61 and genes encoding proteins which activate RhoA (ARHGEF12, GNA11, GNA12, TGFβ1) as well as YAP1 and YES1.

**Figure 4.**
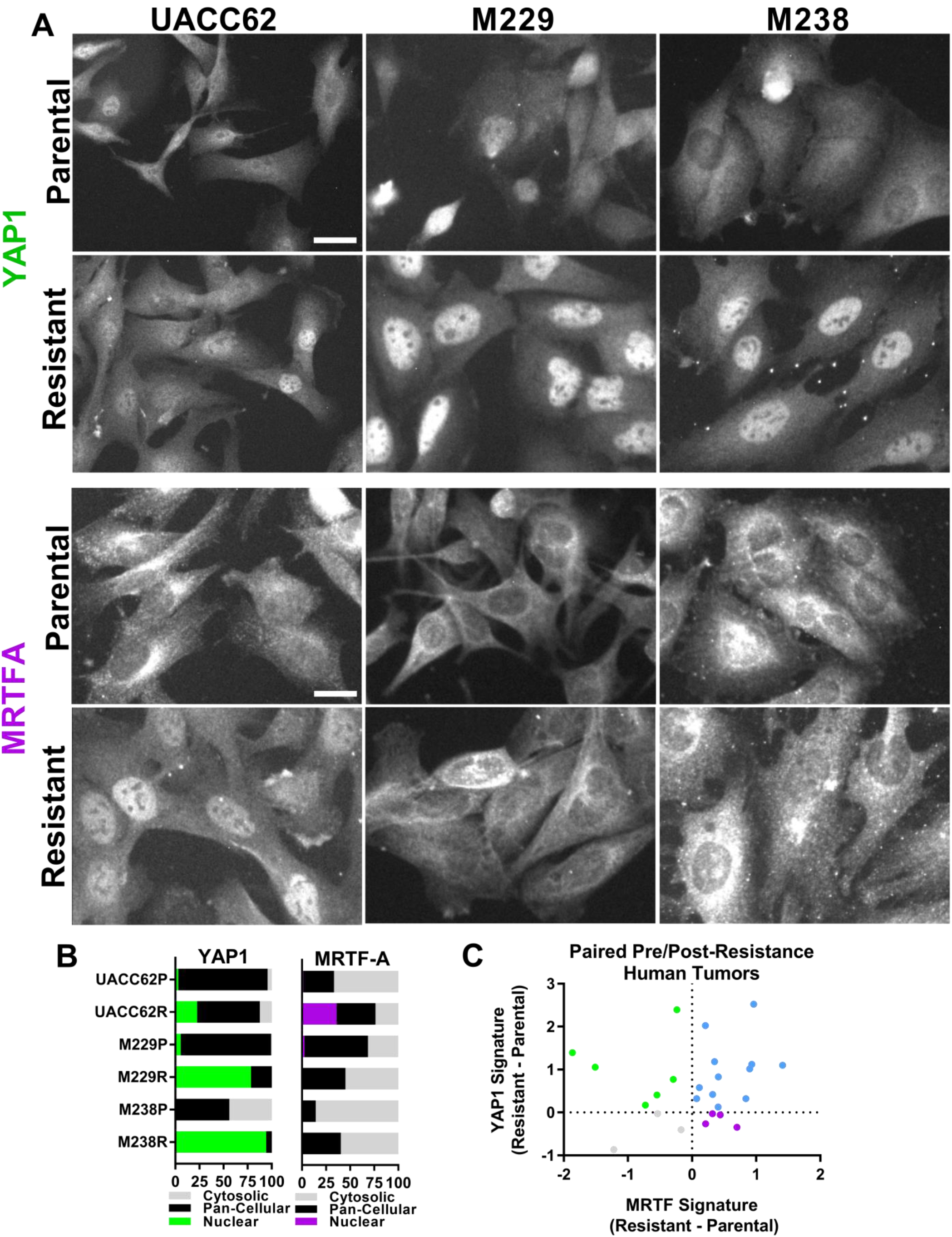
YAP1 and MRTF-A are activated in de-differentiated BRAFi-resistant cells. **A)** M229P/R, M238P/R, and UACC62P/R cells were seeded into 8-well chamber slides and were allowed to attach overnight. The next day, cells were fixed and stained with an anti-YAP1 or anti-MRTF-A antibodies. Representative images from n = 3 biological replicates and n = 1 technical replicates. Scale bar is 5 µm. **B)** Quantification of staining from panel ‘A’. Data are represented as a stacked bar graph wherein the fraction of cells that have predominantly nuclear, pan-cellular, or cytosolic localization is plotted as a fraction of the total cells. **C)** MRTF and YAP1 signatures were predicted for human melanoma tumor pairs which had an increase in RhoA/C signature score from **(Fig. 1G)**. Change in MRTF and YAP1 signature score between baseline and resistant tumors is plotted.

YAP1 and MRTF gene signature activation is increased in the paired pre- and post-resistance human melanoma tumors which had an increase in RhoA/C signature gene expression **(Fig. 4C)**. Out of this subset of tumors, only 3/24 displayed upregulation of neither YAP1 or MRTF target genes. Half (12/24) of the tumors had upregulation of both YAP1 and MRTF gene signatures, which could possibly result from the high degree of redundancy in the transcriptional output from YAP1 and MRTF. Another explanation is that this could result from the tumors consisting of a mixed population of YAP1^High^ and MRTF^High^ cells. Some tumors appeared to have selective activation of YAP or MRTF, which is interesting considering the apparent mutual exclusivity of MRTF-A/YAP1 activation in the experimentally derived cell line models. This is again consistent with the transcriptional alterations in the RhoA^High^ BRAFi-resistant cell lines since MRTFA and YAP1 gene signatures are both increased in the poorly differentiated tumors **(Fig. S10)**. Taken together, these data demonstrate that YAP1 and/or MRTF are activated in nearly all of the poorly differentiated BRAFi-resistant cells/tumors.

### Pharmacologically targeting MRTF/YAP-mediated gene transcription

Since our results indicated that YAP1 and MRTF are activated in de-differentiated BRAFi-resistant cells, we reasoned that pharmacologically targeting these transcriptional mechanisms would be sufficient to re-sensitize cells to vemurafenib. YAP1 is activated by YES1, a Src family kinase. Previous studies have used the Src family kinase inhibitor, dasatinib, to inhibit YES1 and subsequently inhibit YAP1 ^36^ and there is also evidence which suggests that Src itself can activate YAP1 ^20^. Using Src inhibition as an approach to block YAP1 activity is also interesting since our bioinformatics analysis predicted that poorly differentiated human tumors are more sensitive to Src inhibitors, including dasatinib **(Fig. 2F)**. To confirm this in the context of vemurafenib-resistant cells, we treated M229R and M238R cells with dasatinib and measured YAP1 nuclear localization. YAP1’s nuclear localization is decreased in both cell lines upon dasatinib treatment **(Fig. 5A/B)**.

**Figure 5.**
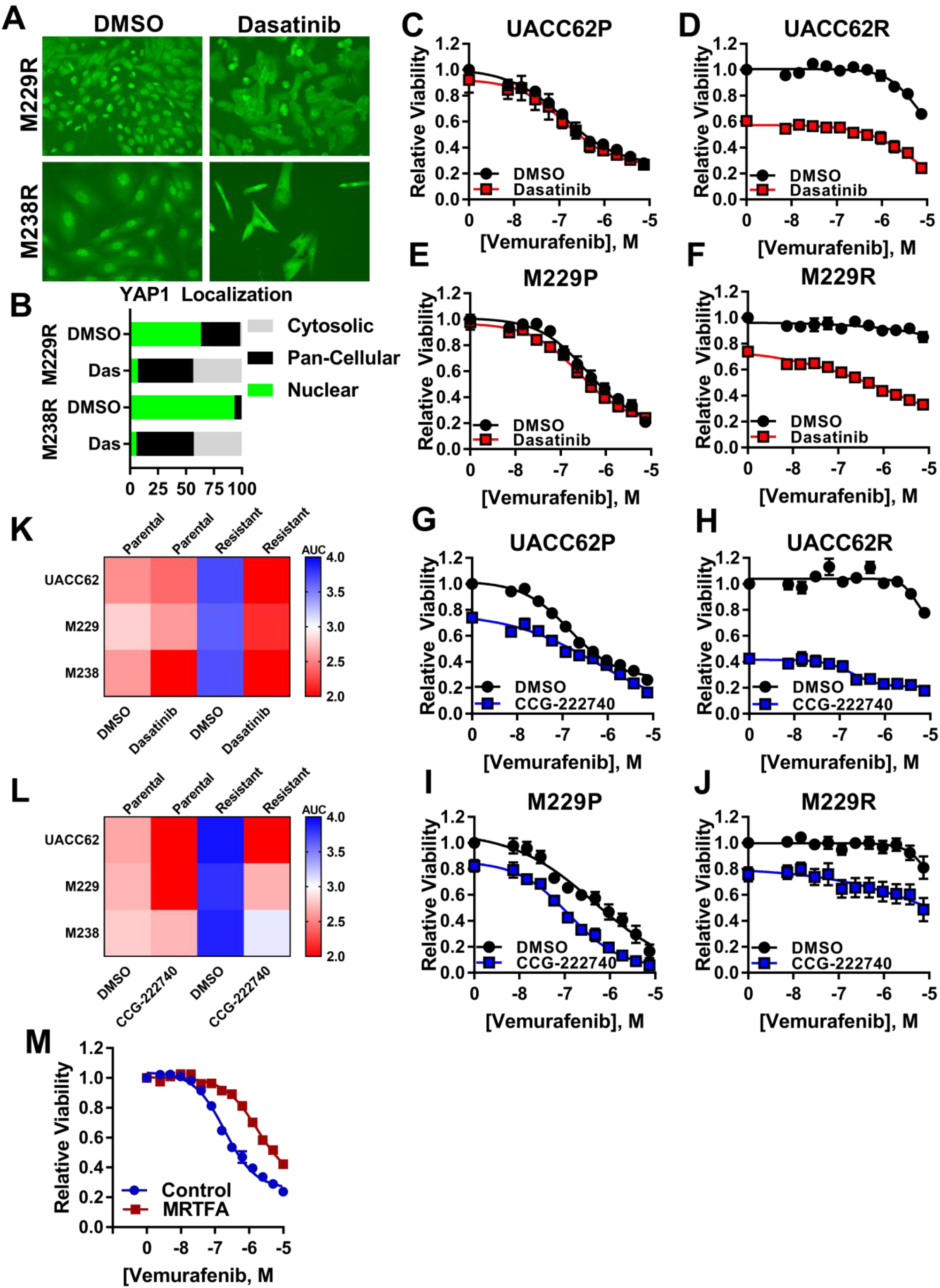
De-differentiated BRAFi-resistant cells are more sensitive to Dasatinib and CCG-222740. **A)** Cells were seeded into 8-well chamber slides and were allowed to attach overnight. The next day, cells were treated with Dasatinib (500 nM) for 16 h, then cells were fixed and stained with an anti-YAP1 antibody. Representative images from n = 3 biological replicates and n = 1 technical replicate. **B)** Quantification of YAP1 localization from panel “A”. Data are represented as a stacked bar graph wherein the fraction of cells that have predominantly nuclear, pan-cellular, or cytosolic localization is plotted as a fraction of the total cells. **C-J)** Parental and Resistant cell lines were seeded into 384-well plates at a density of 1,000 cells/well and cells were allowed to attach overnight. The next day, cells were treated in dose response with Vemurafenib at the indicated concentrations −/+ Dasatinib (red) or CCG-222740 (blue) at 10 µM. At 72 h viability was measured with CellTiter-Glo. Cells and treatments were as indicated with Dasatinib or the MRTF pathway inhibitor CCG-222740. Dose response curves are viability data are from n = 3 biological replicates and n = 1 technical replicate. **K)** Heatmap of AUC values for the Vemurafenib/Dasatinib drug combination for four parental and resistant cell line pairs. **L)** Heatmap of AUC values for the Vemurafenib/CCG-222740 drug combination for four parental and resistant cell line pairs. **M)** UACC62P cells stably expressing Gus (control) or MRTFA were seeded into 384-well plates and treated with a 14-point vemurafenib concentration gradient with a top dose of 10 µM as described in the materials and methods. Data is average from n = 3 biological replicates with n = 3 technical replicates.

We next wanted to determine whether dasatinib re-sensitizes de-differentiated BRAFi-resistant cells to vemurafenib. Dasatinib treatment has only a minor effect on potentiating the vemurafenib response in the parental UACC62P and M229P cells **(Fig. 5C/E)**, however, the vemurafenib response is greatly potentiated in the resistant UACC62R and M229R cells **(Fig. 5D/F)**. While UACC62R does not have as robust YAP1 activation as M229R and M238R, the minor increase in YAP1 nuclear localization could explain why these cells also respond to dasatinib. This effect is consistent across all three de-differentiated BRAFi-resistant cell lines **(Fig. 5K)**. All three of the de-differentiated BRAFi-resistant cell lines also have increased sensitivity to dastatinib as a single agent **(Fig. S11)**.

Our lab has developed a series of MRTF pathway inhibitors, including CCG-222740 ^1, 10, 15, 23^ **(Fig. S12)**. We sought to determine whether this inhibitor can re-sensitize de-differentiated BRAFi-resistant cells to Vemurafenib. CCG-22740 has only a modest effect on re-sensitizing M229R or M238R cells, which have strong YAP1 but low MRTF-A activation **(Fig. 5I/J/L)**. CCG-222740 has the strongest re-sensitization effect in UACC62R cells **(Fig. 5G/H/L)** which was the only BRAFi-resistant cell line with strong nuclear localization of MRTF-A. Also, UACC62R cells are more sensitive to CCG-222740 as a single agent **(Fig. S11)**. To more directly determine the effect of MRTF-A on BRAFi resistance we generated cells which stably express wildtype MRTF-A. Cells expressing MRTF-A are approximately 10-fold less sensitive to vemurafenib **(Fig. 5M)**. Interestingly when we performed the inverse experiment, deletion of MRTF-A in resistant cells did not alter vemurafenib sensitivity **(Fig. S13)**. Although we did not observe any overt change in MRTF-B localization when parental and resistant cell lines were compared under basal conditions **(Fig. S14)**, it is possible that MRTF-A depletion may induce MRTF-B activation. Or given the functional redundancy between MRTF and YAP1, it is possible that MRTF-A depletion could induce compensatory YAP1 activation. Taken together these data demonstrate that inhibition of RhoA-mediated gene transcription in de-differentiated melanoma cells, which can be mediated either by YAP1 or MRTF, re-sensitizes the melanoma cells to vemurafenib.

## Discussion

In this study we sought to identify a pharmacological “Achilles heel” for BRAFi-resistant melanoma cells/tumors. In theory, if pathway-centric dependences can be identified for cells with acquired resistance, then co-targeting these resistance pathways concurrently with MAPK pathway inhibitors may delay, prevent, or reverse resistance. We found evidence for RhoA pathway activation in approximately half of BRAFi/MEKi-resistant human melanoma cells and tumors. In isogenic BRAFi-resistant cell lines, Rho pathway activation was accompanied by both an increase in actin stress fibers and usually MLC2 phosphorylation. These findings are consistent with previous reports which demonstrate that actin stress fibers are increased in cell line models of acquired BRAFi resistance ^17^. Building off these findings, we demonstrated that ROCK inhibition re-sensitizes Rho^High^ BRAFi-resistant cells to vemurafenib, highlighting the importance of this signaling pathway in adaptive BRAFi resistance. This finding also supports our bioinformatics predictions, since multiple ROCK inhibitors were among the drugs predicted to be selective for poorly differentiated melanoma tumors.

We next wanted to identify signaling mechanisms which are associated with RhoA pathway activation. These signaling mechanisms could serve as biomarkers for RhoA activation or these pathways could directly promote RhoA activation. Upon acquisition of drug resistance all of the RhoA^High^ downregulate an array of melanocyte lineage genes such as TYR, MLANA, and SOX10. This is accompanied by upregulation of multiple cancer invasion-associated genes including AXL and SOX9 as well as several collagen and integrin isoforms. De-differentiation of melanoma cells has previously been linked to drug resistance. For instance, a switch in MITF/AXL gene expression levels mark BRAFi resistance ^19, 46, 49^. In another study silencing of SOX10, which was one of the most downregulated genes in our analysis, promotes BRAFi resistance ^48^. But whether de-differentiation is directly inducing RhoA activation, or if RhoA activation is simply associated with de-differentiation is a question that still needs to be addressed.

As a result of modulating the actin cytoskeleton, Rho regulates gene transcription. Rho-induced F-actin polymerization allows for MRTF and YAP1 to translocate into the nucleus where they subsequently regulate gene transcription ^5, 26, 27, 33, 39, 44^. Interestingly, some reports suggest that MRTF and YAP1 physically interact and are present in close proximity on similar gene promoters ^56^, while others suggest more indirect mechanisms of shared gene expression control ^8^. While YAP1 has been previously demonstrated to promote BRAFi resistance in melanoma ^7, 14, 17, 22^, the role of MRTF in BRAFi resistance is unknown. In this study we demonstrate that nuclear accumulation of either MRTF-A or YAP1 is increased in RhoA^High^ BRAFi-resistant cells. We also demonstrate that pharmacologically inhibiting MRTF-mediated transcription increases vemurafenib sensitivity. Conversely, we also demonstrate that overexpression of MRTF-A induces vemurafenib resistance. Further work is required to determine the nature by which MRTF promotes BRAFi resistance and whether those signaling mechanisms are similar to the mechanisms by which YAP1 promotes BRAFi resistance. Interestingly, we observed YAP1 activation in 2 of 3 RhoA^High^ resistant cell lines, and MRTF-A activation in the 3^rd^ cell line. This may suggest that MRTF-A and YAP1 are acting redundantly in this context and that activation of either MRTF-A or YAP1 is sufficient to promote drug resistance. It will also be important to determine whether deletion of MRTF-A induces compensatory activation of other MRTF isoforms or perhaps YAP1 since MRTF-A deletion has no effect on vemurafenib sensitivity.

This study demonstrates that Rho^High^ BRAFi-resistant cells are re-sensitized to vemurafenib by ROCK inhibitors and that this Rho^High^ phenotype is linked to de-differentiation. The direct signaling mechanisms which lead to Rho activation in melanoma cells are still unclear, but it is enticing to suggest that induction of TGFβ upon Sox10 loss ^48^ may lead to RhoA activation. However, it is possible that TGFβ may be inducing de-differentiation ^30, 54^ and RhoA activation simultaneously through different signaling mechanisms. Future studies will be necessary to elucidate details of these signaling networks. While it is already known that YAP1 promotes BRAFi resistance, these studies build upon that knowledge to demonstrate that dasatinib blocks the nuclear accumulation of YAP1 and enhanced drug sensitivity in BRAFi-resistant cells. Since dasatinib is already FDA-approved for other indications, it highlights the potential of a re-purposing approach for treatment of BRAFi/MEKi-resistant melanomas. In this context, dasatinib may be most effective in combination with vemurafenib since in at least one resistant cell line vemurafenib potency was restored to that of parental cells. These studies also link MRTF-A activation to BRAFi resistance for the first time, highlighting the potential of targeting MRTF-mediated transcription to prevent or treat drug resistant melanoma. In total, these studies provide robust predictions of precision therapy approaches to prevent or treat clinical BRAFi resistance based on pharmacological inhibition of RhoA-mediated gene transcription.

