## Supplementary figures and images for "Rho-mediated gene transcription promotes BRAF inhibitor resistance in de-differentiated melanoma cells"

### Figure S1

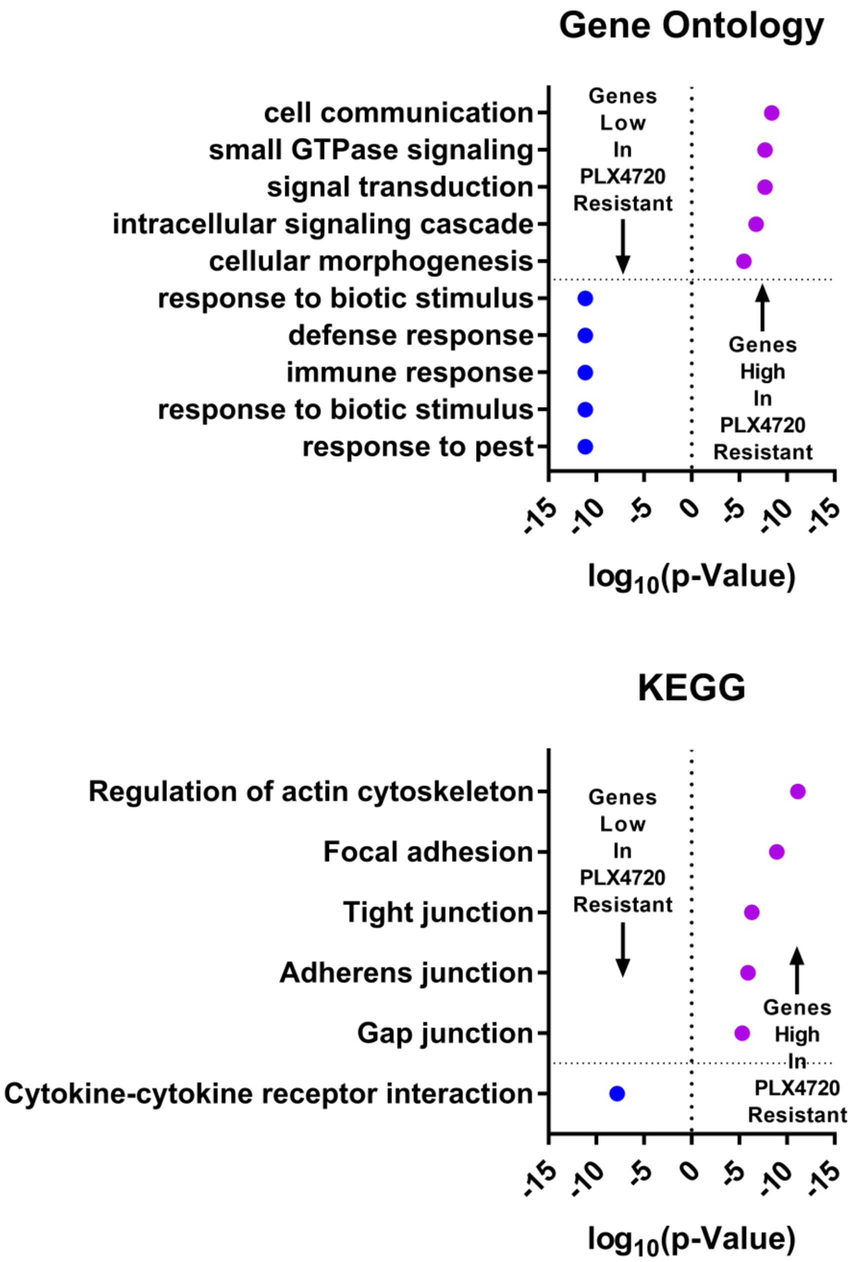

### Figure S2

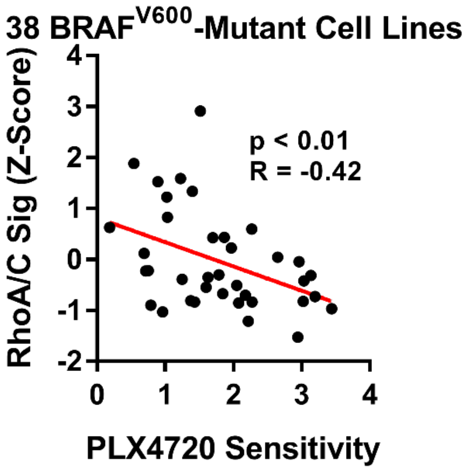

### Figure S3

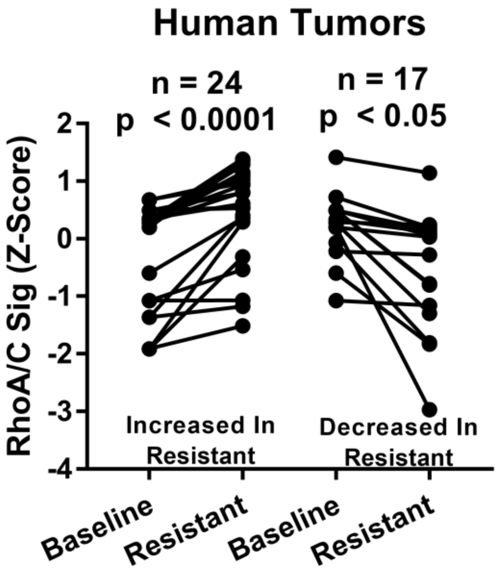

### Figure S4

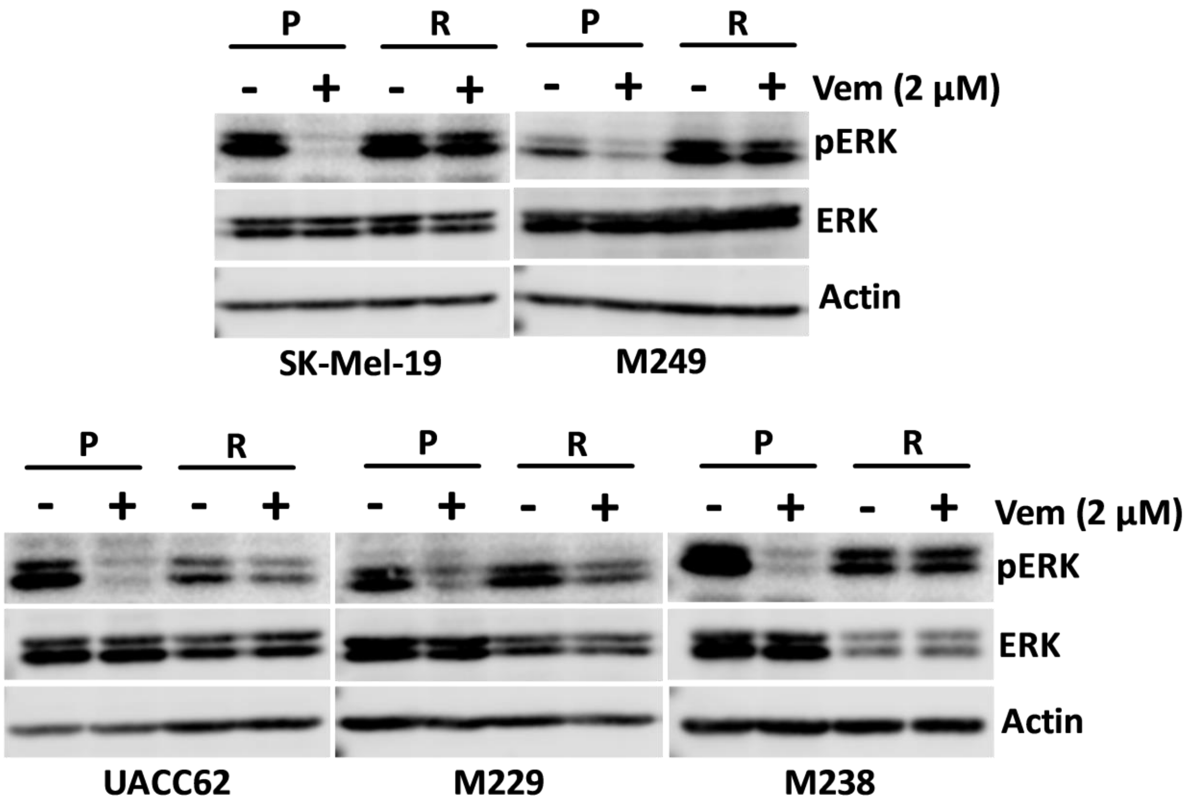

### Figure S5

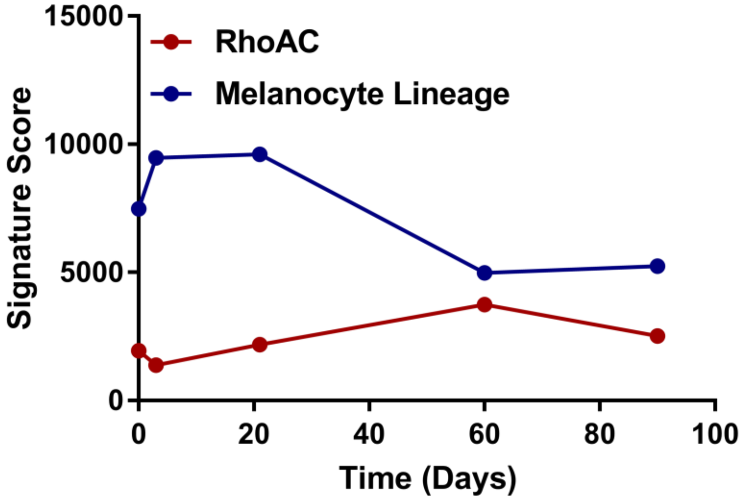

### Figure S6

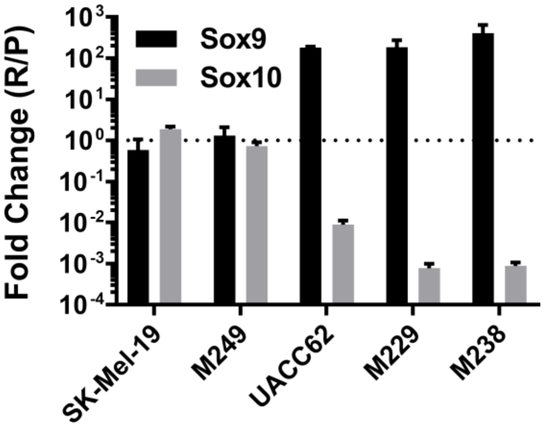

### Figure S7

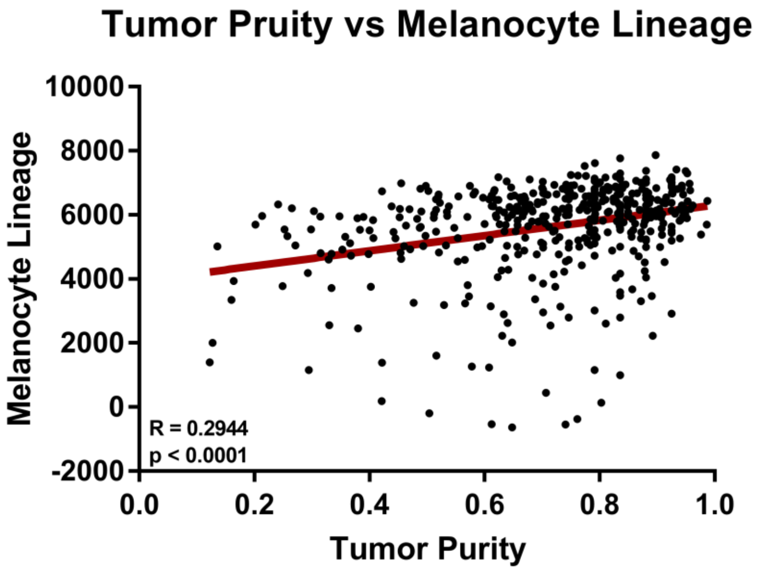

### Figure S8

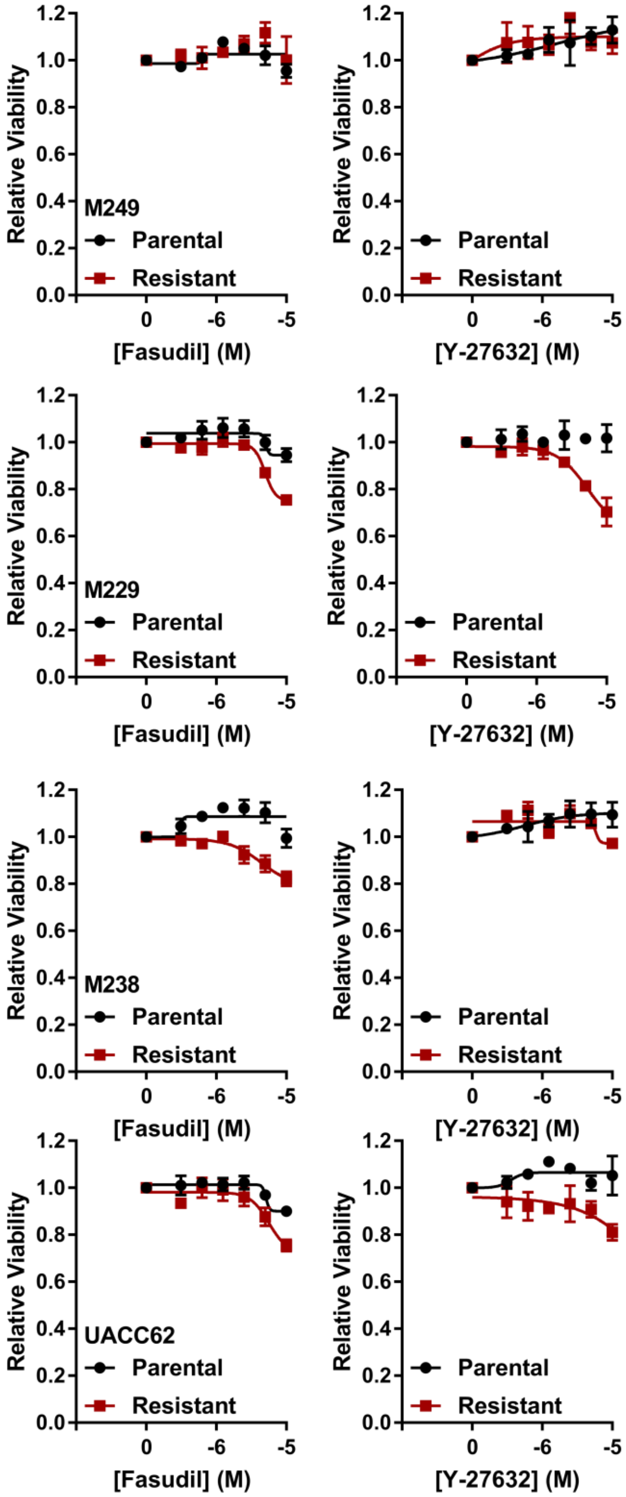

### Figure S9

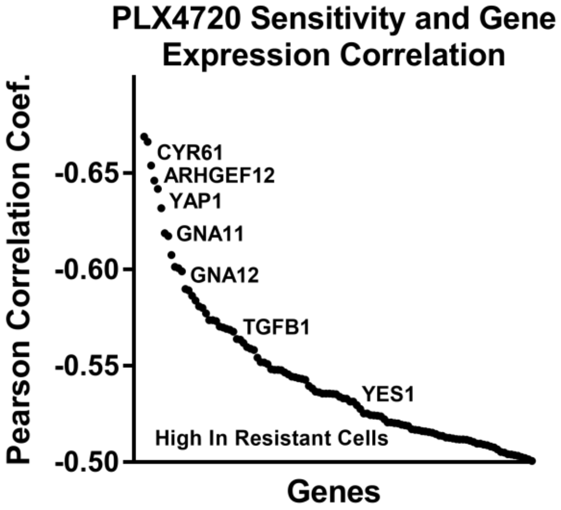

### Figure S10

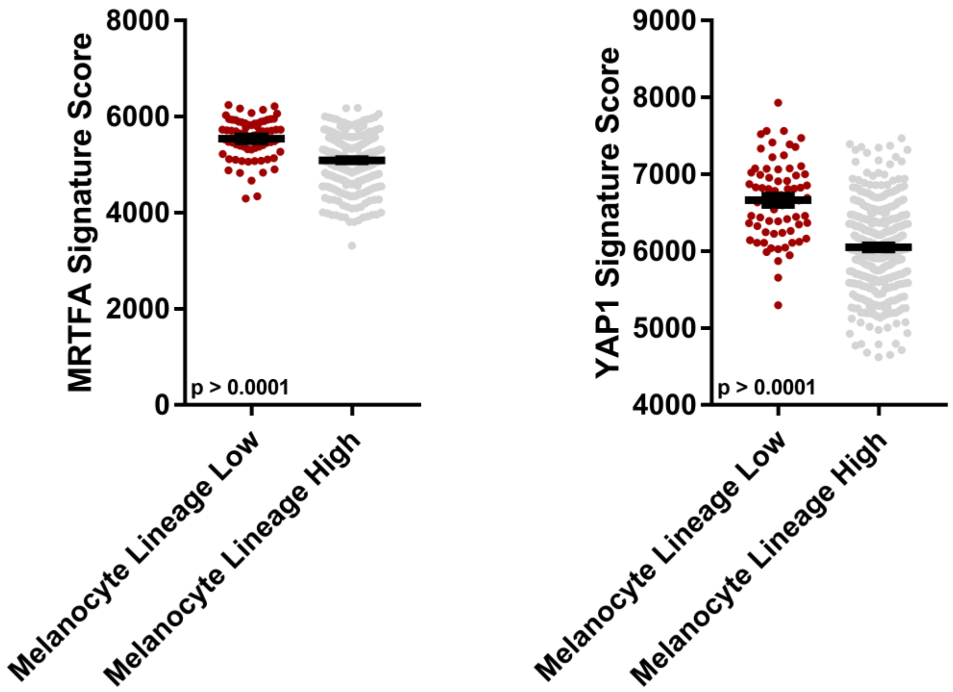

### Figure S11

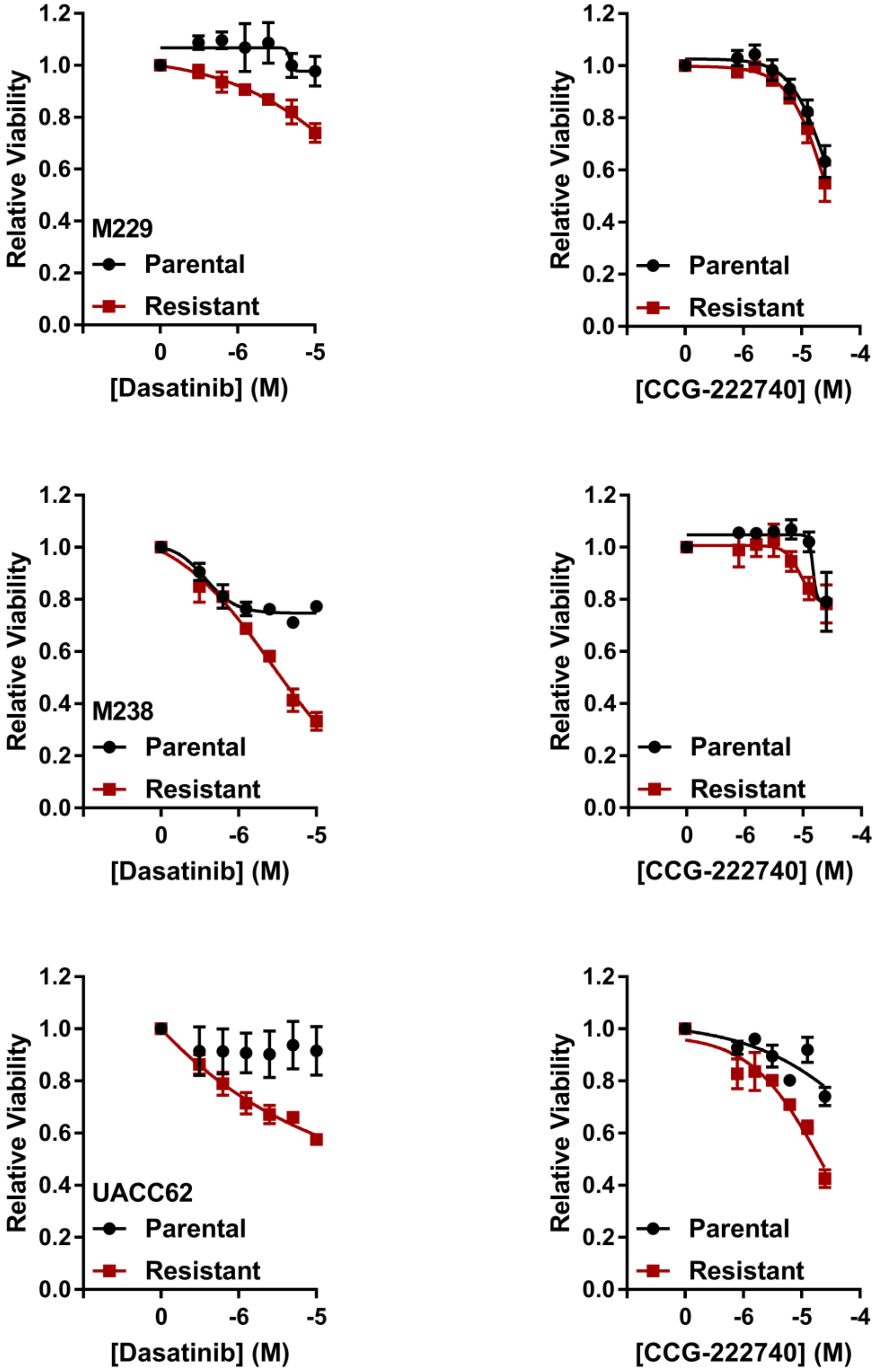

### Figure S12

**Figure S12**

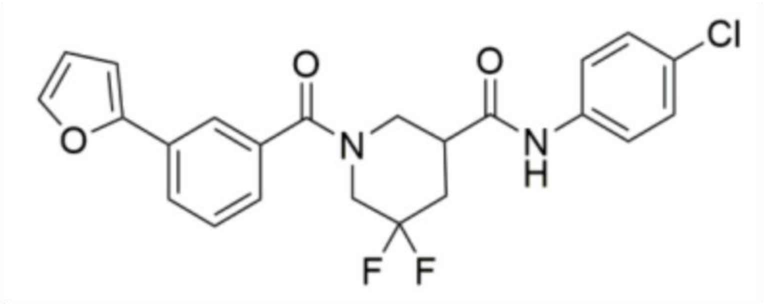

### Figure S13

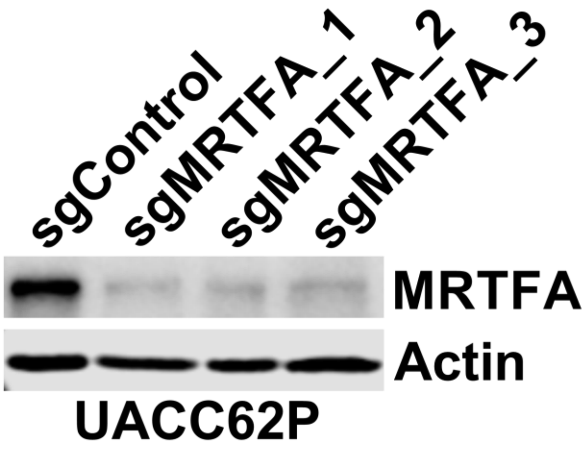

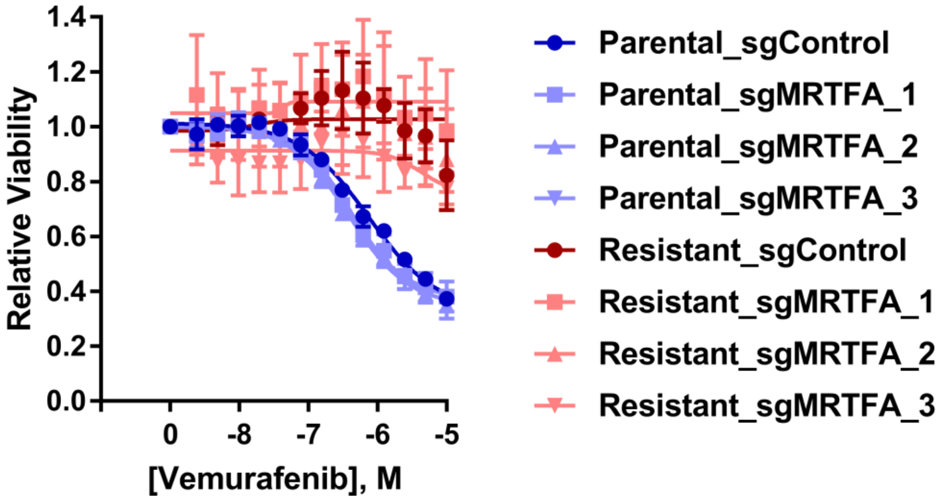

### Figure S14

**Figure S14**

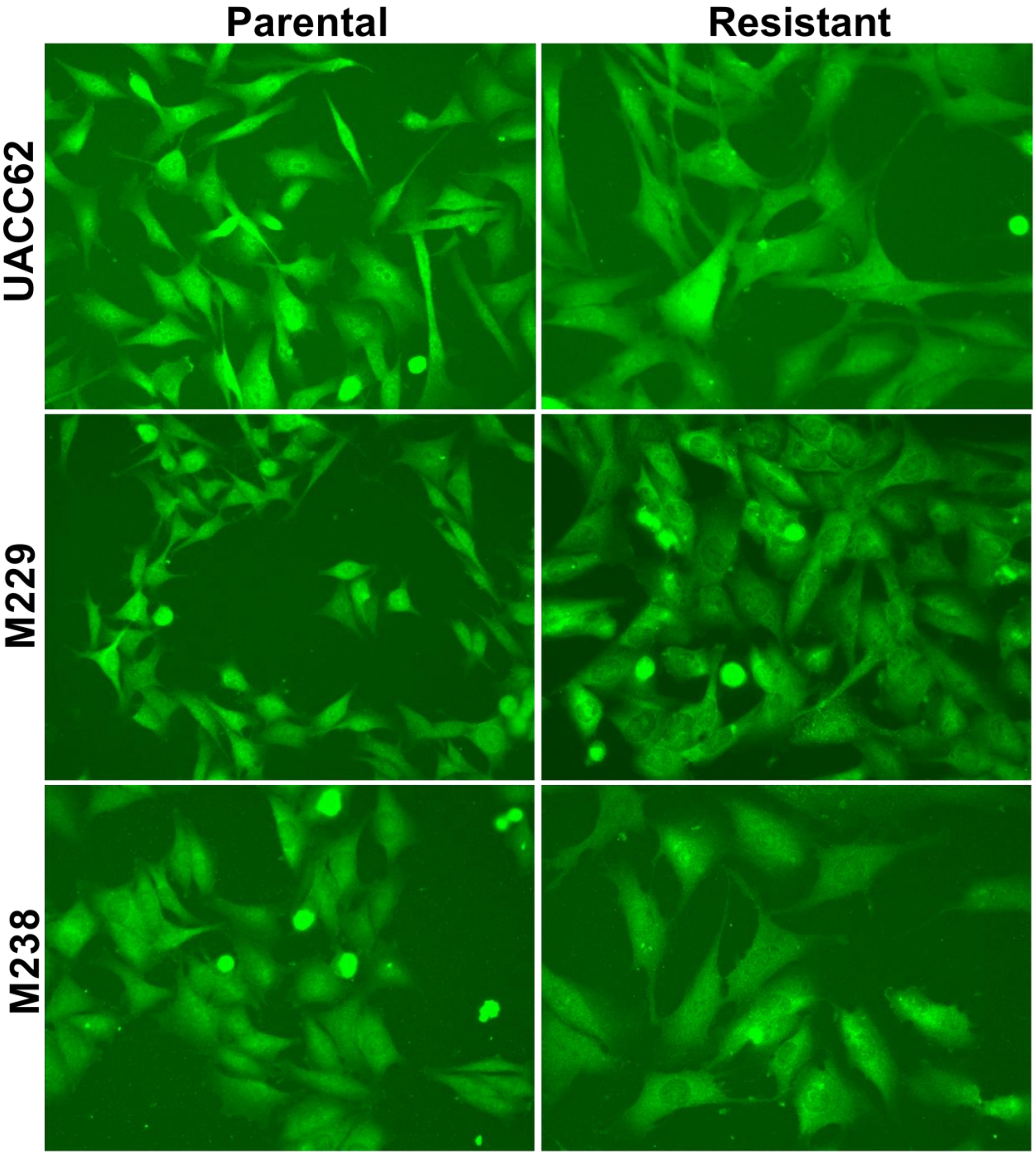
